# The status of current and tentative marine natural World Heritage areas

**DOI:** 10.1101/2021.02.28.433275

**Authors:** Caitlin D. Kuempel, B. Alexander Simmons, Madeline Davey

## Abstract

The 1972 World Heritage Convention (WHC), along with the 1994 Global Strategy, aim to preserve the outstanding universal value of internationally important cultural and natural sites within a “representative, balanced and credible” network of highly-protected areas. Increasing human pressures and shortfalls in representation have been documented across the World Heritage network, particularly in terrestrial and cultural sites, threatening the integrity and primary goals of the WHC. However, the conservation status of current and tentative (i.e., proposed) marine natural World Heritage areas remains relatively unknown. We assessed the extent of recent (2013) and historical (2008-2013) cumulative human impacts and several metrics of representation (country, continent, ecoregion, wilderness, and threatened species) within existing and tentative marine natural World Heritage areas. We found moderate yet increasing cumulative human impacts across most existing sites, and high or very high impacts across the majority of tentative sites. Climate change impacts comprised nearly 75% of impact scores, on average, and differences in land and marine impacts across sites could help prioritise management decisions. Over 75% of marine ecoregions and 85% of threatened species considered in this study have no representation within the existing marine natural World Heritage network. We outline examples of how prioritizing representation across tentative sites for future World Heritage listing could greatly increase these measures. We urge the WHC to adopt quantitative, systematic and transparent evaluations of how current and tentative sites contribute to the overarching goals of maintaining a representative World Heritage network and preserving their outstanding universal values for future generations.

## Introduction

The 1972 World Heritage Convention (WHC) was designed to safeguard some of Earth’s most extraordinary natural and cultural sites from being lost (UNESCO 1972) and is one of the most significant international environmental agreements to date. Today, 193 state parties have committed to “ensuring the identification, protection, conservation, presentation and transmission to future generations of the cultural and natural heritage” for the 1,100+ properties that have been deemed to have outstanding universal value to humanity as a whole (UNESCO 1972). World Heritage areas (WHAs) are considered the highest level of recognition afforded globally (UNESCO 2019) and contribute to international targets aimed to benefit both people and nature, such as the Convention on Biological Diversity Aichi Target 11 (Secretariat of the CBD 2010) and UN Sustainable Development Goals 11.4, 14, and 15 (UN General Assembly 2015).

The primary goal of the WHC is to recognize and maintain the outstanding universal value of listed sites in perpetuity (UNESCO 1972). To meet outstanding universal value standards, a natural site must meet at least one of the ten selection criteria (Table 1), the conditions of integrity (intactness and wholeness), and have adequate management and protection to maintain outstanding universal value in the future (UNESCO 2019). Given there are practically no areas free from human influence (Halpern et al. 2015; Jones et al. 2018), quantifying the status of threats within current and tentative natural WHAs is integral to meeting and maintaining outstanding universal value standards. Recent research uncovered potential shortfalls in the terrestrial natural WHA estate, documenting an average 63% increase in human pressures (Allan et al. 2017), potentially compromising the outstanding universal value of several sites. Increasing human pressures have also been documented within marine natural WHAs (mnWHA), including a 22% decline in coral reef fish abundance per year in the Ningaloo Marine Park WHA (Vanderklift et al. 2019) and a loss of around 50% of coral within the Great Barrier Reef WHA (De’ath et al. 2012). The human impacts driving these declines, however, have yet to be quantified across the global mnWHA estate.

**Table 1.** Selection criteria for inclusion on the World Heritage List based on outstanding universal value of cultural and natural protected areas. Beyond these ten criteria, the protection, management, authenticity and integrity of properties are also important considerations for the World Heritage Convention (UNESCO 2019).

| **Selection criteria** |  | **Type of property** |
| --- | --- | --- |
| i. | Represent a masterpiece of human creative genius | Cultural |
| ii. | Exhibit an important interchange of human values, over a span of time or within a cultural area of the world, on developments in architecture or technology, monumental arts, town-planning or landscape design | Cultural |
| iii. | Bear a unique or at least exceptional testimony to a cultural tradition or to a civilization which is living or which has disappeared | Cultural |
| iv. | An outstanding example of a type of building, architectural or technological ensemble or landscape which illustrates (a) significant stage(s) in human history | Cultural |
| v. | An outstanding example of a traditional human settlement, land-use, or sea-use which is representative of a culture (or cultures), or human interaction with the environment especially when it has become vulnerable under the impact of irreversible change | Cultural |
| vi. | Directly or tangibly associated with events or living traditions, with ideas, or with beliefs, with artistic and literary works of outstanding universal significance. (The Committee considers that this criterion should preferably be used in conjunction with other criteria) | Cultural |
| vii. | Contain superlative natural phenomena or areas of exceptional natural beauty and aesthetic importance | Natural |
| viii. | Outstanding examples representing major stages of earth's history, including the record of life, significant on-going geological processes in the development of landforms, or significant geomorphic or physiographic feature | Natural |
| ix. | Outstanding examples representing significant on-going ecological and biological processes in the evolution and development of terrestrial, fresh water, coastal and marine ecosystems and communities of plants and animals | Natural |
| x. | Contain the most important and significant natural habitats for in-situ conservation of biological diversity, including those containing threatened species of outstanding universal value from the point of view of science or conservation | Natural |

In 1994, secondary objectives of creating a “representative, balanced and credible” World Heritage List (UNESCO 1994a) were implemented to address imbalances of WHAs, particularly within cultural sites and across regions. However, imbalances in WHA representation are known to be widespread: cultural sites have higher representation than natural sites, natural sites are biased towards terrestrial areas, and biases exist in geographic locations and species coverage (UNESCO 1994a; IUCN 2004; Abdulla et al. 2014). Notably, there is no explicit definition or guideline as to what constitutes a “representative, balanced and credible” WHA network, making it difficult to evaluate and measure progress toward achieving these goals.

Sites of global significance must be safeguarded and their outstanding universal value maintained while capturing a more comprehensive range of natural values within the World Heritage List. Here, we use best available data to assess current mnWHAs in terms of representation and cumulative human impacts to provide an updated reference point to measure progress toward representation (Abdulla et al. 2014) and a baseline for managing impacts within mnWHAs. We then assess how the list of tentative mnWHAs complement existing sites to more efficiently achieve progress towards these goals in the future.

## Methods

### World heritage areas

Spatial boundaries of existing WHAs were obtained from the World Database on Protected Areas (WDPA; UNEP-WCMC & IUCN 2020). We consider existing mnWHAs as those established as of January 2020. We only included natural or mixed cultural/natural properties designated as either marine or mixed terrestrial/marine. Terrestrial areas were excluded from mixed sites for analysis. WHAs with less than 10 km^2^ of marine area were excluded to maximise the reliability of averaged impact estimates, resulting in the exclusion of five sites. A total of 43 existing mnWHAs were included in the analysis. Tentative WHAs were identified by comparing the most recent tentative list (World Heritage Centre 2019) with the equivalent boundary within the WDPA (see Text S1 and S2 for details). Following the same approach described above, we included 63 tentative mnWHAs in the analysis. All data for existing and tentative mnWHAs included in the analysis are available in Tables S1 and S2, respectively.

### Representation within marine natural World Heritage areas

We assessed the 43 existing and 63 tentative mnWHAs in terms of the representation of rare ecoregions, wilderness, and threatened species. Data on ecoregions included the global extent of 232 marine ecoregions (coast and shelf areas) (Spalding et al. 2007) and 37 pelagic provinces (surface waters) (Spalding et al. 2012). Marine ecoregions are a biogeographic classification of broad-scale patterns of marine species and communities, and are commonly used to measure representation in systematic conservation planning (Butchart et al. 2015; Kuempel et al. 2016; Jantke et al. 2018).

Measures of ecoregion rarity, wilderness, and threatened species were chosen due to their direct relationship with the selection criteria for outstanding universal value (UNESCO 2019). Ecoregion rarity speaks to the evolving interpretation of outstanding universal value as incorporating both uniqueness and representativeness (World Heritage Committee 1996). We define the rarest ecoregions as the smallest 20% of ecoregions based on global extent. Wilderness is defined as “biologically and ecologically intact seascapes that are mostly free of human disturbance” (Jones et al. 2018) and relates to selection criteria vii, ix and x for aesthetic importance, ecological and biological significance and, arguably, for its significance for *in-situ* conservation of biological diversity (UNESCO 2019). The global extent of marine wilderness was obtained from Jones et al. (2018) and is classified as areas representing the bottom 10% of the cumulative human impact and its individual stressors. Threatened species representation relates to selection criteria x of the WHC. Spatial data for species ranges were obtained from the IUCN Red List of Threatened Species databases (IUCN 2017). We focus on species that may utilise the existing habitat(s) across all mnWHAs, thus excluding plants and reef-forming corals that define habitat characteristics unique to each site. Freshwater fishes, molluscs, and dragonflies/damselflies were excluded due to a lack of comprehensive data, and birds were excluded for their exceptionally large geographic ranges on average. We also excluded any species classified as ‘data deficient’ or ‘least concern’, and any species ranges that are known to be extinct. Subspecies were aggregated at the species level for reporting. We calculated the number of threatened species ranges within each site, as well as a threatened species density relative to the size of the WHA’s marine area.

### Cumulative human impacts

We used the most recent map (2013) of marine cumulative human impacts (CHI) (Halpern et al. 2015). CHI represents a continuous scale of human impact at a 1 km^2^ resolution based upon 19 stressors. We divided stressors into three categories: climate-based (indirect human impacts), land-based and ocean-based (direct human impacts). Climate-based pressures include sea surface temperature, UV radiation, ocean acidification, and sea level rise. Ocean-based stressors include fishing (destructive fishing, non-destructive low bycatch, non-destructive high bycatch, pelagic low bycatch, pelagic high bycatch, and artisanal fishing), shipping routes, invasive species, and ocean pollution. Land-based stressors include population density, night-lights, fertilizer pollution, pesticide pollution, and other inorganic pollution. We excluded oil rigs from our calculations due to their lack of impact within mnWHAs.

Changes in CHI (ΔCHI) from 2008 to 2013 were estimated using data from Halpern et al. (2015). Data for sea level rise, shipping routes, invasive species, and ocean pollution were not available for 2008 and were excluded from calculations. Ocean acidification, artisanal fishing, and inorganic pollution were also excluded because they did not differ between 2008 and 2013. We calculated the average CHI and ΔCHI within the marine boundaries of existing and tentative sites. Similarly, we calculated the average impact for each of the 19 pressures (stressor intensity × habitat vulnerability). We divided the global CHI 2013 data into five quantiles of impact: < 2.358 (*very low*), 2.358–3.325 (*low*), 3.325–3.749 (*moderate*), 3.749–4.172 (*high*), > 4.172 (*very high*).

## Results

### Existing marine world heritage areas

#### Representation

The existing mnWHA network consists of approx. 1.39 million km^2^ of marine area across 48 sites, representing 0.34% of global marine area (Fig. 1a). Of the 43 existing mnWHA sites included in this study, most (51%) are not exclusively marine but contain some terrestrial area, with an average of 53% of marine area within each site. By number, the majority of mnWHAs are located in Oceania (23%) and Europe (23%), followed by Asia (20%), North America (18%), Africa (9%), and South America (7%) (Fig. 1b). By country, Australia contains the greatest number of mnWHAs (6 sites), with Costa Rica, France, Indonesia, the United Kingdom, and the United States each containing 2 sites. Oceania also contains the biggest mnWHAs by area (58%; 802,415 km^2^), followed by North America (28%) and South America (11%). Europe contains as many sites as Oceania, but these sites only cover 2% of the total WHA estate. Sites within Africa and Asia contribute less than 1% each.

**Figure 1.**
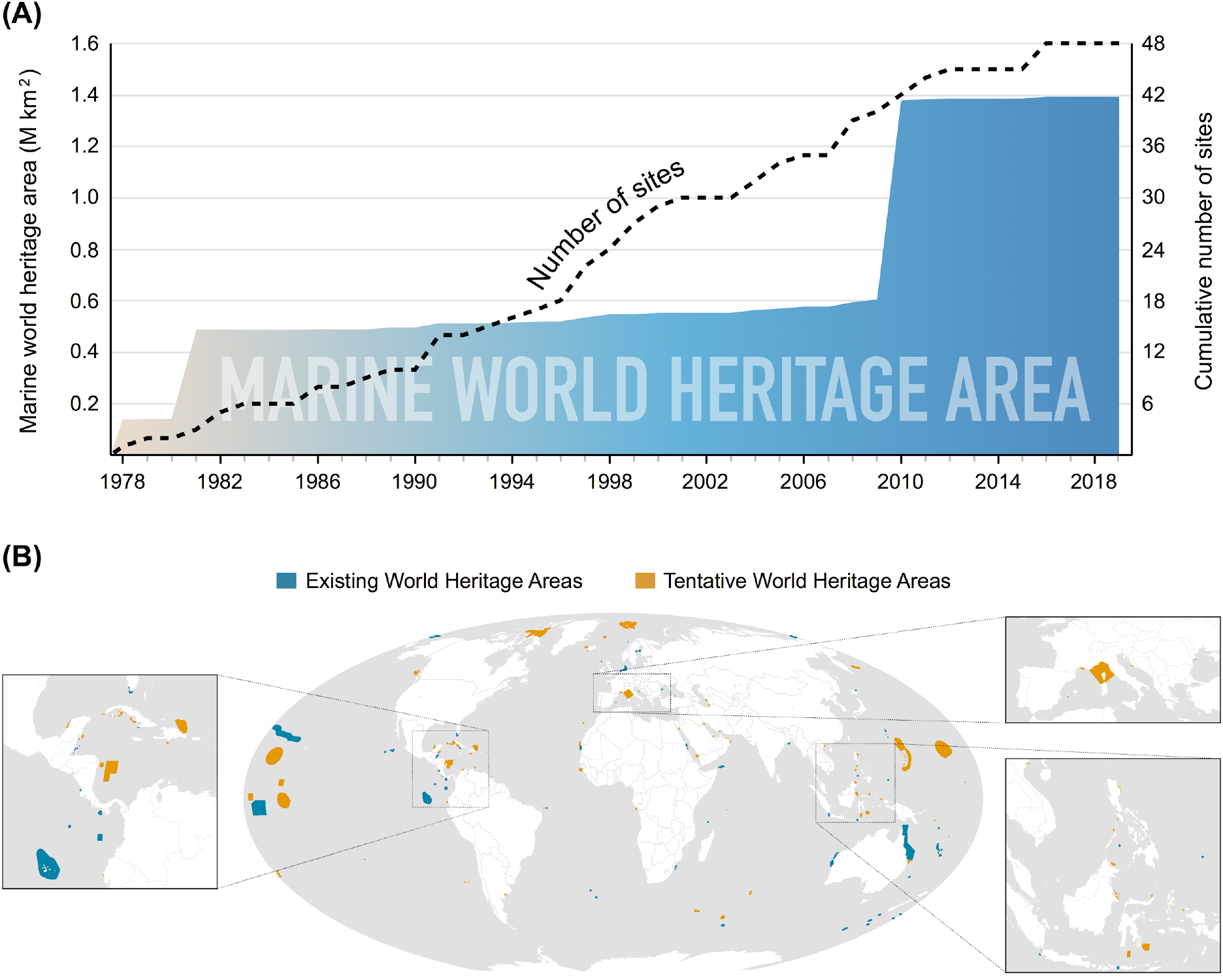
(a) The cumulative number and extent of marine world heritage areas over time. (b) Spatial extent of existing and tentative marine world heritage areas as of 2019. For mixed terrestrial/marine sites, only the respective marine extent is shown. Site boundaries have been thickened to enhance visibility.

Only 4% (54,632 km^2^) of the mnWHA network consists of marine wilderness, which constitutes roughly 0.11% of the global marine wilderness area. Nearly half of all sites contain some marine wilderness (44%), but the area covered within each site is only 15%, on average. In Oceania, mnWHAs are 5.82% marine wilderness, and the fraction of wilderness in mnWHAs in all other regions is less than 1%. Asia does not contain any marine wilderness within mnWHAs and Europe has only 0.02%. Sites with the greatest proportion of marine wilderness include Macquarie Island, Wrangel Island Reserve, New Zealand Sub-Antarctic Islands, and Heard and McDonald Islands (Table S1).

The current mnWHA network includes 24% of the planet’s marine ecoregions and 35% of all pelagic provinces. Most of these marine ecoregions are only represented within one mnWHA, with the exception of the Western Mediterranean and Shark Bay ecoregions (two sites each). The pelagic provinces with greatest representation include the Eastern Tropical Pacific (five sites), North Central Pacific Gyre (three sites), and Non-Gyral Southwest Pacific (three sites). Thirteen (30%) existing mnWHAs contain the rarest marine ecoregions in the world, including those of the Cocos Islands, Galapagos Islands, and Fernando de Naronha and Atol das Rocas.

The mnWHA network overlaps with the ranges of 402 (13%) threatened species. On average, each site overlaps with 53 threatened species ranges, with a density of 0.162 species km^−2^. The majority of threatened species within these sites are classified as ‘vulnerable’ (40%), followed by ‘near threatened’ (28%), ‘endangered’ (22%), and ‘critically endangered’ species (10%). Chondrichthyes (sharks, rays, and chimaeras) represent the greatest proportion of threatened species within the mnWHA network (59%), followed by marine fish, marine mammals, reptiles, and sea cucumbers. Collectively, cone snails, amphibians, terrestrial mammals, and freshwater shrimp, crayfish, and crabs represent 2% of all threatened species. Sites with the greatest density of threatened species include the Gulf of Porto, Ogasawara Islands, Surtsey, and Ibiza (Table S1).

#### Cumulative human impacts

Existing mnWHAs are experiencing moderate impacts, with a global average cumulative human impact (CHI) of 3.52 ± 0.84 in 2013. Ten sites (23%) exhibit high impacts (CHI > 3.75), and nine sites (21%) exhibit very high impacts (CHI > 4.17) (Fig. 2a). Only 37% of sites exhibit low or very low impacts (CHI < 3.33). The greatest impacts were observed in mnWHAs throughout Asia (CHI = 3.983), followed by Europe, Africa, North America, South America, and Oceania (CHI = 2.963) (Fig. 2b). Sites experiencing the greatest CHI include Rock Islands Southern Lagoon, Ujung Kulon National Park, Komodo National Park, and the Sundarbans (Table S1). Climate-based impacts contributed to 73% of all CHI scores on average. Ocean-based impacts contributed to 20% of CHI, with 8% attributed to fishing, 7% to shipping and invasive species, and 5% to ocean pollution. Land-based impacts contributed to less than 8% of CHI across all mnWHAs. Only a few sites were primarily impacted by these direct impacts, such as Ibiza, Ha Long Bay, the Sundarbans, and the Wadden Sea (Table S1).

**Figure 2.**
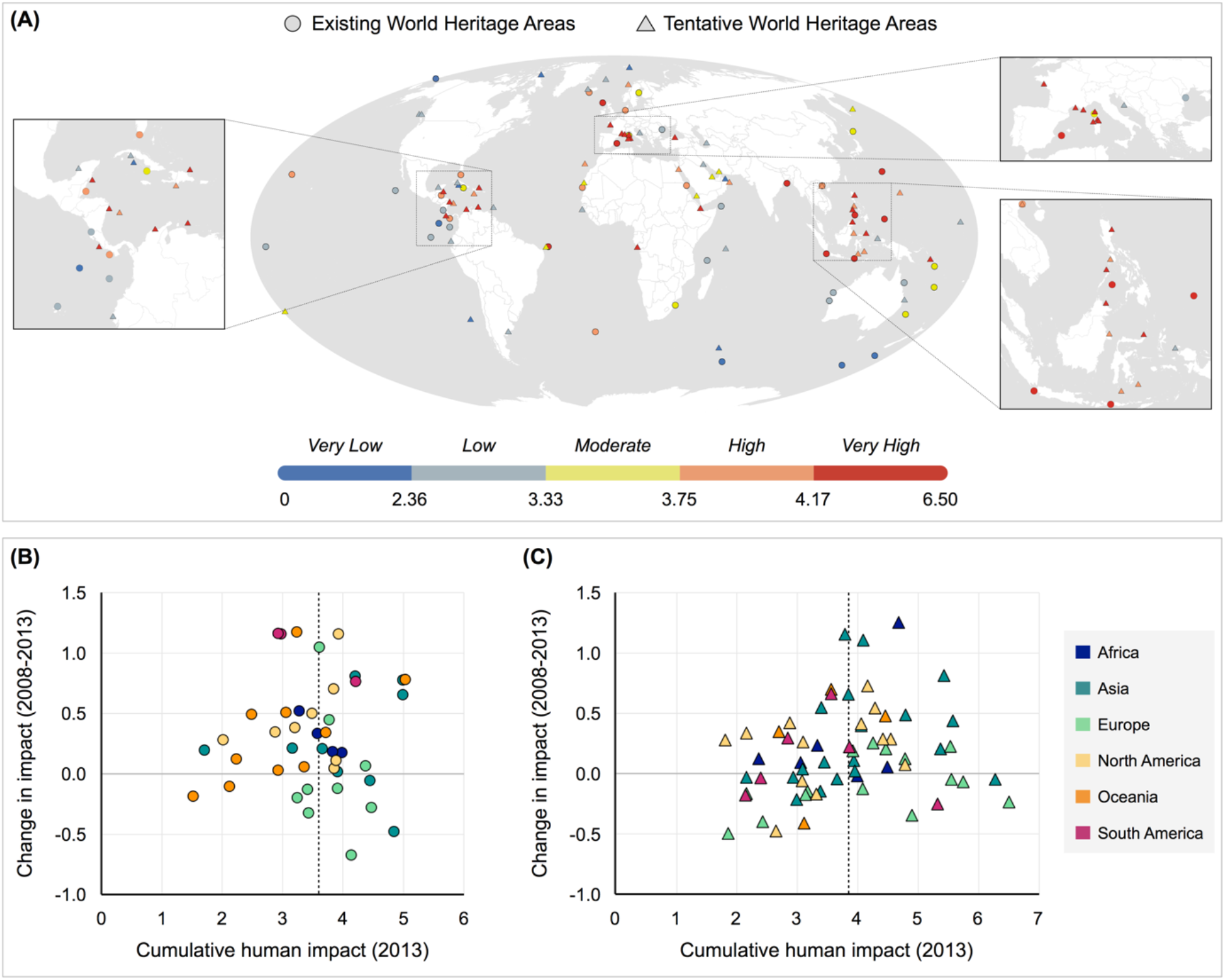
(a) Average cumulative human impact (CHI) of existing and tentative marine world heritage areas as of 2013. Relationship between average CHI (2013) and the average change in CHI since 2008 for (b) existing and (c) tentative marine world heritage areas. Dotted line represents the median cumulative human impact.

The majority of existing mnWHAs (77%) experienced an increase in CHI between 2008 and 2013 (Fig. 2b). Fourteen sites (33%) experienced increases greater than 0.50, and five sites (12%) experienced increases greater than one. One site, Surtsey, experienced a reduction in CHI greater than 0.50. The greatest increases occurred in South American mnWHAs (ΔCHI = +1.032), followed by North America, Oceania, Africa, Asia, and Europe (ΔCHI = −0.015). Sites experiencing the greatest increase in CHI since 2008 include Ningaloo Coast, Galapagos Islands, Coiba National Park, and Malpelo Fauna and Flora Sanctuary (Table S1). Despite relatively high impacts in 2013, sites like the Sundarbans, St. Kilda, and Surtsey experienced decreasing impacts since 2008. Climate-based impacts exhibited the greatest average increase across all sites (ΔCHI = +0.348). Overall, ocean- and land-based impacts have exhibited minimal change since 2008, with average ocean-based impacts slightly decreasing ((ΔCHI = − 0.040) and land-based impacts remaining relatively unchanged (ΔCHI = +0.003).

### Tentative marine natural World Heritage areas

#### Representation

Of the 63 tentative mnWHAs, 29% are located in Asia, 27% in Europe, 20% in North America, 9% in South America, 9% in Africa, and 6% in Oceania (Fig. 2a). Countries with the greatest number of proposed mnWHAs include France, Indonesia, the Philippines, Canada, Italy, and Norway (Table S2). Nearly all tentative mnWHA sites include terrestrial areas (92%). A much larger proportion of the tentative mnWHA network includes marine wilderness compared to the existing network (18%; 366,810 km^2^), which is spread across 15 sites (24%) (Table 2a).

**Table 2.**
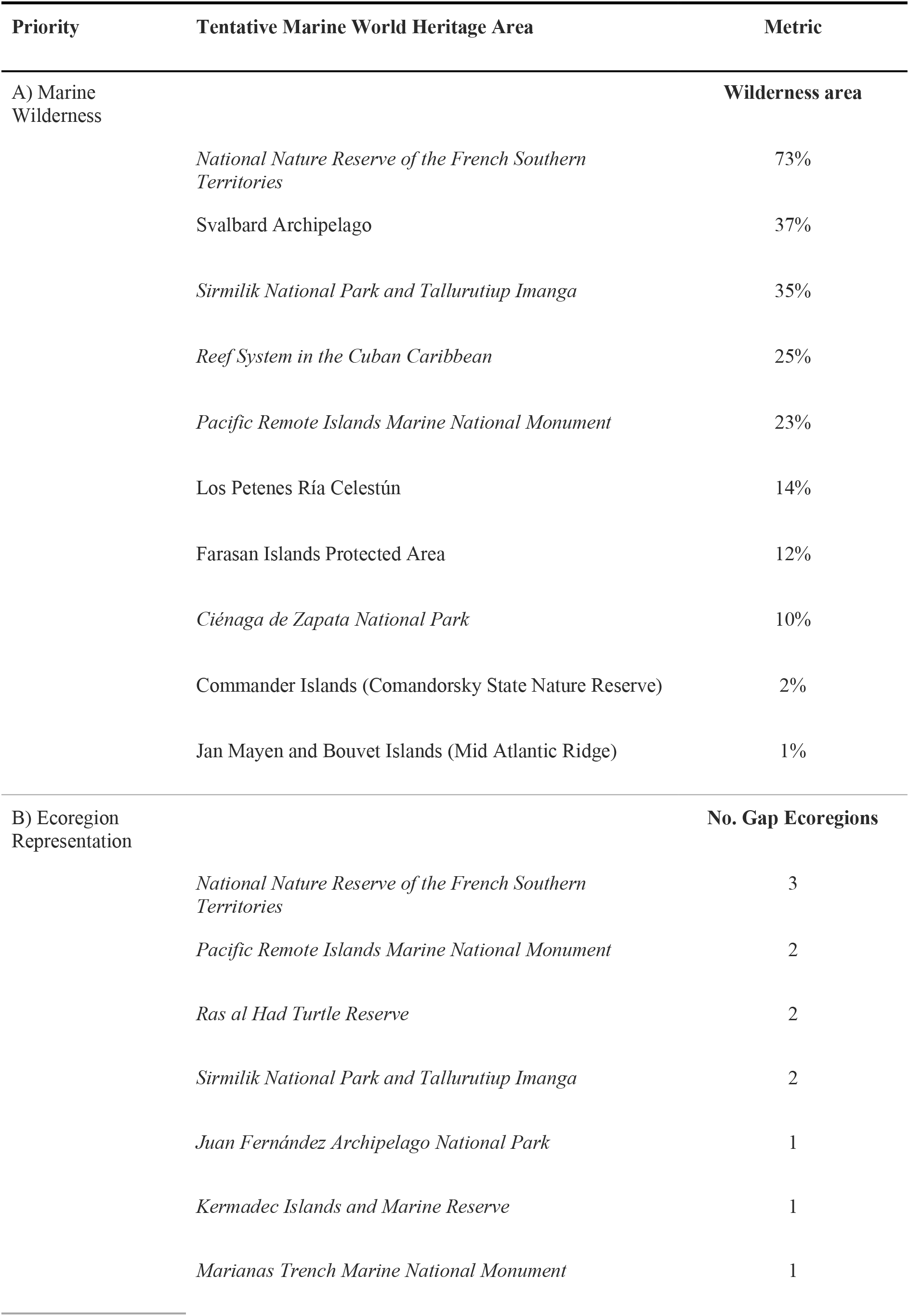
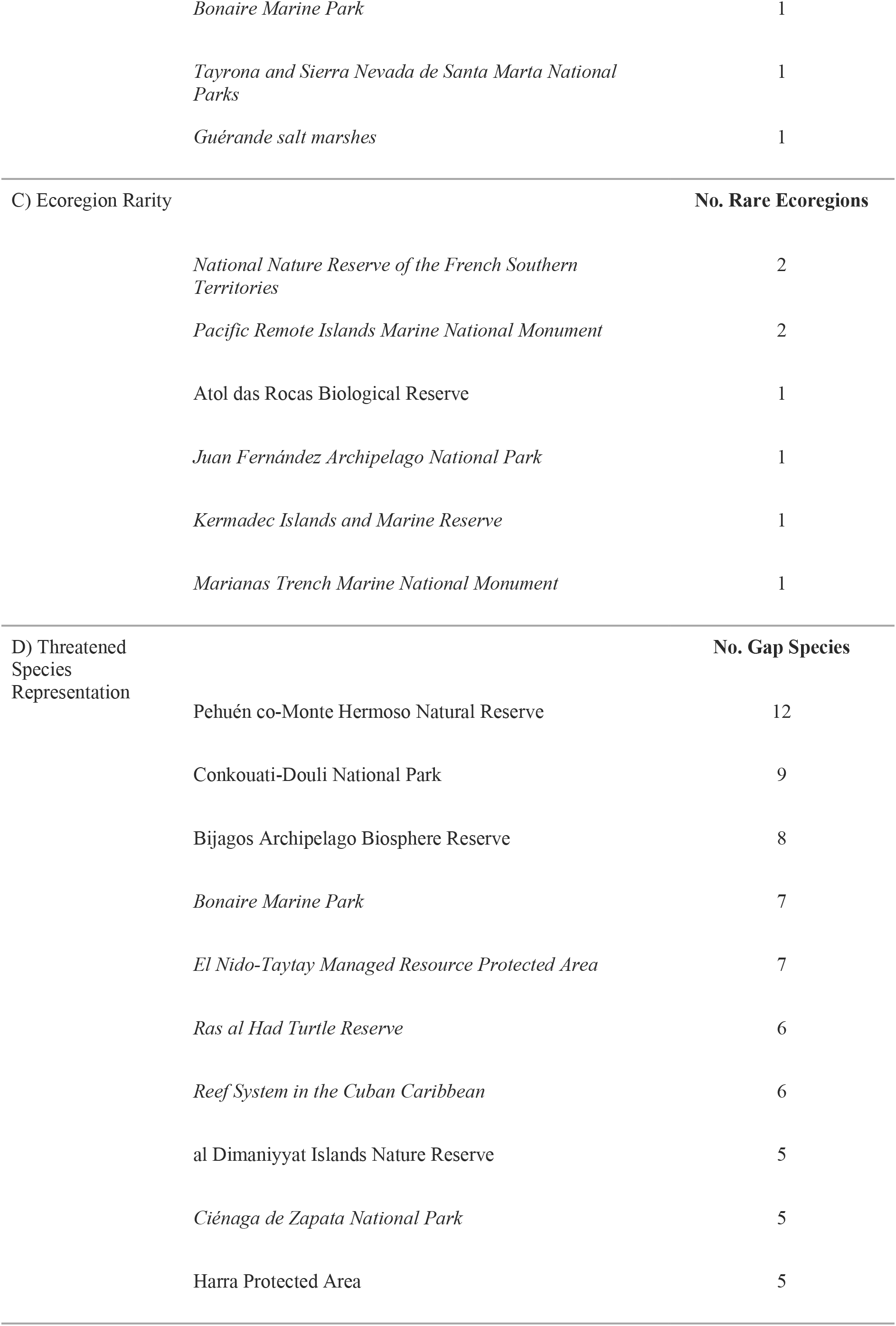
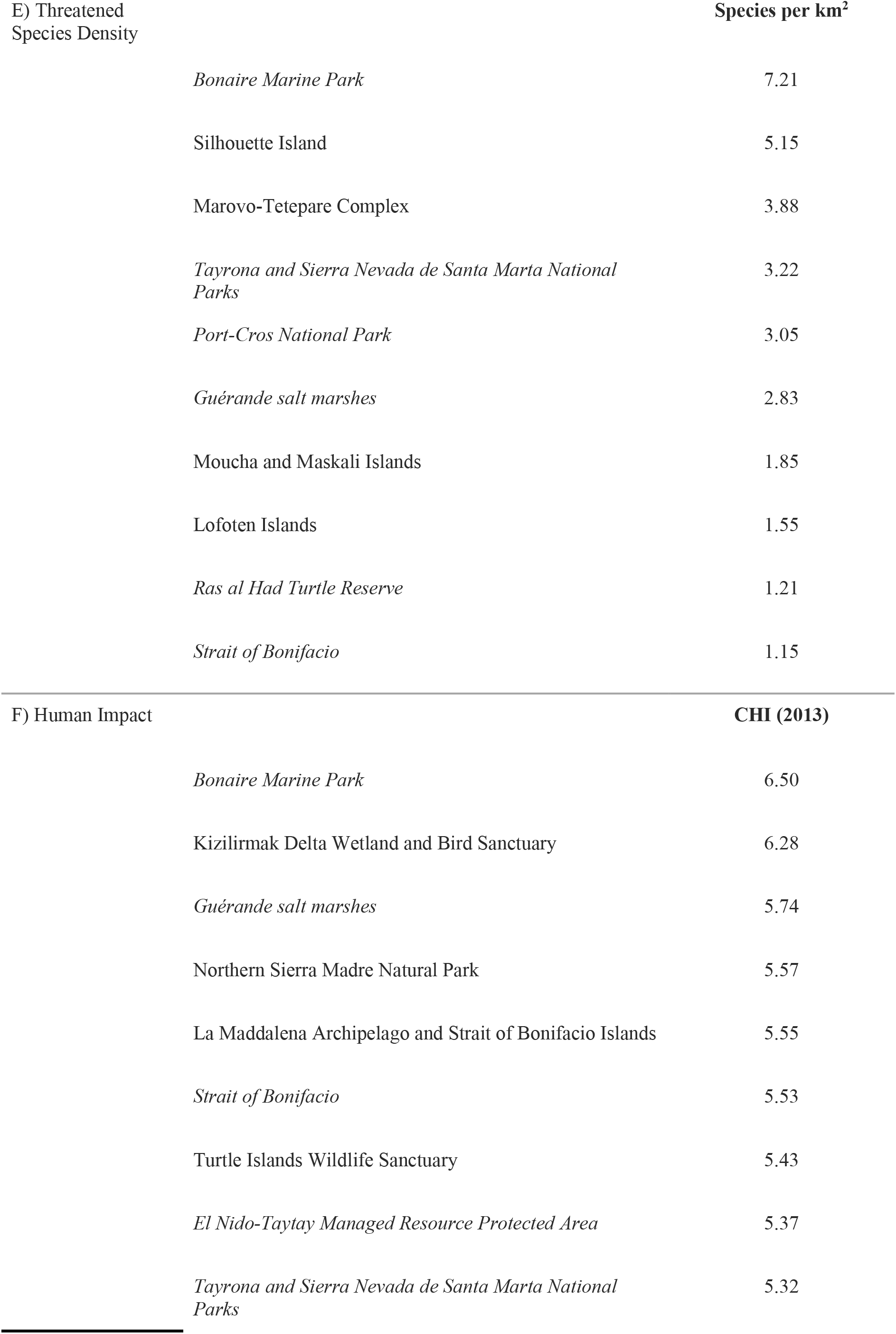
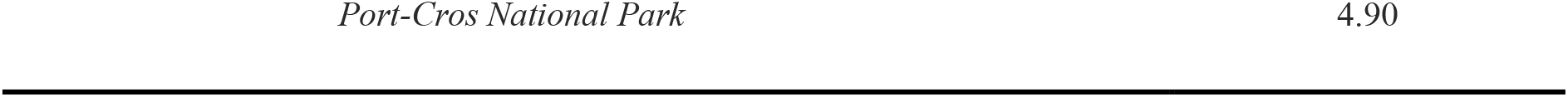
Top 10 tentative marine nature World Heritage areas (mnWHAs) under different priorities for maximising representation, uniqueness, and conservation. Sites that are present under multiple priorities are in *italics*.

Thirty-seven (69%) marine ecoregions within tentative mnWHAs are not currently represented in the existing mnWHA network (Table 2b). Six (16%) of these ‘gap’ ecoregions constitute the rarest marine ecoregions globally, such as the Amsterdam-Saint Paul, Juan Fernández and Desventuradas, and Kermadec Island ecoregions (Table 2c). Further, 50% of pelagic provinces within tentative sites are unrepresented within existing mnWHA, including the Subarctic Pacific, Arctic, Inter American Seas, and North Central Atlantic Gyre.

Tentative mnWHAs overlap with 368 threatened species ranges, of which 39% are classified as ‘vulnerable,’ 29% ‘near threatened,’ 21% ‘endangered,’ and 11% ‘critically endangered.’ Although the average number of species per site is similar to that of existing mnWHAs (55 spp.), the average density per site is nearly four times greater (0.600 spp. km^−2^) (Fig. 3). Like existing mnWHAs, Chondrichthyes make up the greatest proportion of threatened species ranges in tentative mnWHAs (62%), followed by marine fish, marine mammals, reptiles, and sea cucumbers. Cone snails, freshwater species, amphibians, and terrestrial mammals collectively make up 3% of species (Table S2). A total of 93 species ranges (25%) are not covered by existing mnWHAs, including 14 (4%) endangered and three (1%) critically endangered species—the Brazilian guitarfish (*Pseudobatos horkelii*), striped smoothhound (*Mustelus fasciatus*), and slender-snouted crocodile (*Mecistops cataphractus*). Tentative mnWHAs with the most unrepresented threatened species and highest density of threatened species can be found in Table 2d,e.

**Figure 3.**
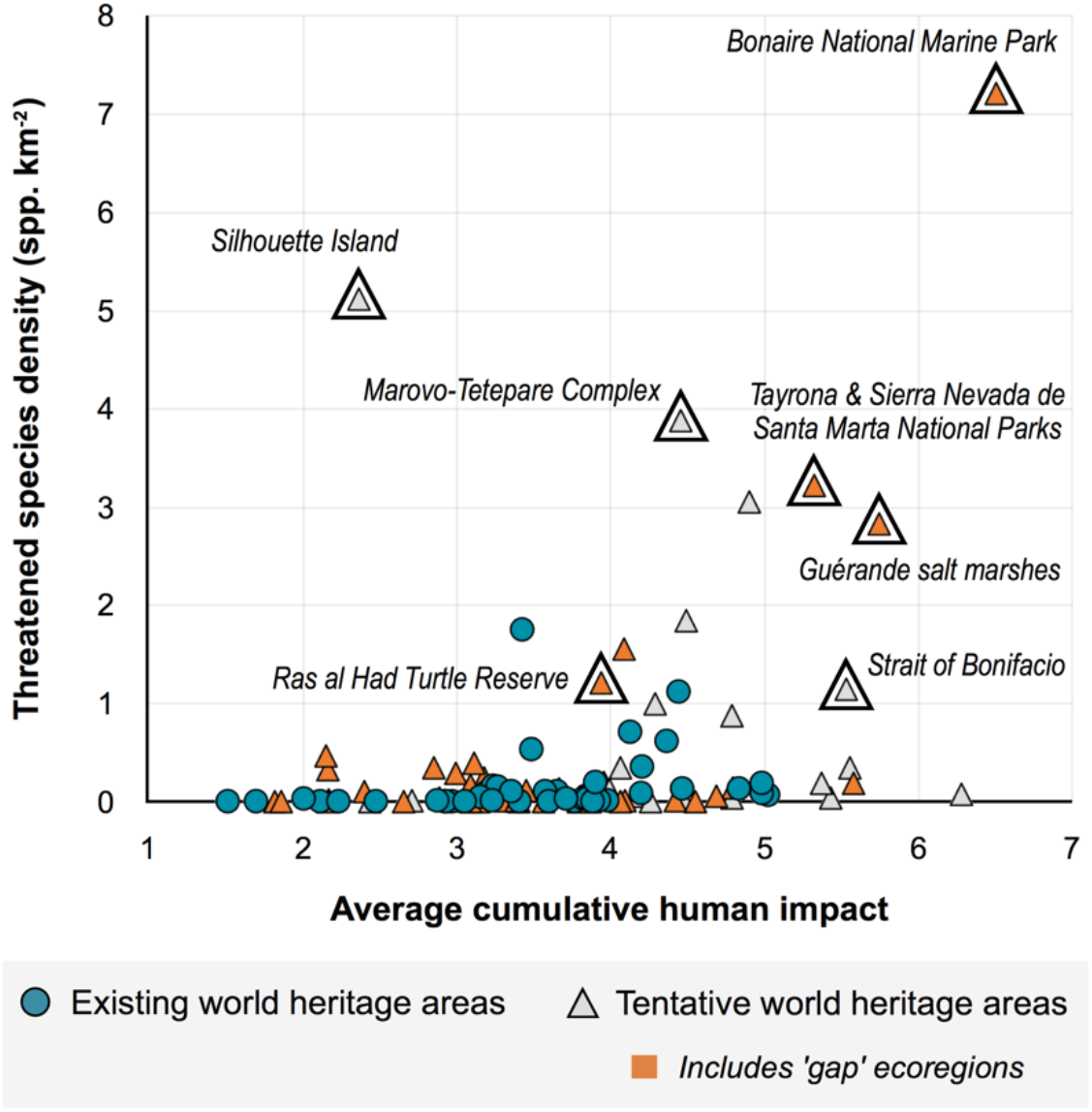
Characteristics of existing and tentative marine world heritage areas reflecting current human impacts, threatened species density, and marine ecoregion representation. Identified sites represent tentative world heritage areas that are of potential significance and are mentioned frequently in the text.

#### Cumulative human impacts

The majority of tentative mnWHAs are under significant threat, with an average CHI of 3.81 ± 1.11 in 2013. Twelve sites (19%) exhibit high impacts, and 21 sites (33%) exhibit very high impacts (Fig. 2a). Bonaire National Marine Park and Kizilirmak Delta Wetland and Bird Sanctuary experience the highest impacts (CHI > 6.0) (Table 2f). Only 37% of tentative sites exhibit low or very low impacts. On average, tentative mnWHAs within Europe are experiencing the greatest human impact (CHI = 4.166), followed by Asia, Africa, North America, Oceania, and South America (Fig. 2c). Climate-based impacts pose the greatest threat to the majority of tentative sites, contributing to 66% of CHI, on average. Ocean-based impacts contribute to 24% of CHI, and land-based impacts contribute nearly 10% (Table S2).

While most tentative mnWHAs have increased CHI since 2008 (62%), changes tend to be much smaller than within existing mnWHAs (Fig. 2c). Only ten sites (16%) experienced increases greater than 0.5, with three sites (5%) showing increases greater than one. The site experiencing the greatest reduction in impact was the National Nature Reserve of the French Southern Territories (ΔCHI = −0.495). Overall, continent-level changes are much smaller than existing sites. The greatest impacts occurred in Asia (ΔCHI = +0.290), followed by Africa, Oceania, North America, South America, and Europe (ΔCHI = −0.082) (Fig. 2c). Like the existing mnWHA, climate-based impacts exhibited the greatest increase across all sites, though at much smaller magnitude (ΔCHI = +0.103). Direct ocean- and land-based impacts have remained relatively unchanged on average (ΔCHI = −0.004).

## Discussion

Our analyses quantified the state of current mnWHAs to identify gaps in achieving WHC goals, including maintaining outstanding universal value and creating a representative WHA network. Though one of the most important conservation agreements, the WHC has not been applied to its fullest potential for mnWHAs. We revealed shortfalls in representation within the current network across all considered measures (geographic location, ecoregions, wilderness, threatened species) and moderate yet increasing cumulative human impacts across the majority of sites, which could undermine conservation efforts. This imperative baseline allowed us to identify tentative sites that could best close these gaps. Our work underscores the need for clear, quantifiable metrics and targets for WHC objectives and the transparent evaluation and reporting of how newly-inscribed WHAs contribute to overarching goals and global biodiversity targets. Although an area’s outstanding universal value is the key requirement within the WHC, it is still important to achieve greater representation as part of the WHC global strategy to reflect distinct ecological values that are underrepresented at this high level of protection (Badman et al. 2008).

### Closing representation gaps

Our results found similar biases in mnWHA representation across continents and countries as recognized previously for all WHAs, like the lack of representation in Africa (UNESCO 1994b, 1994a). Country considerations are important for nomination and management (UNESCO 2019), but we suggest that the WHC take steps to implement additional priorities based on ecologically meaningful criteria—such as wilderness, ecoregions, and threatened species—to ensure greater representation, balance and credibility across natural sites that contribute to conservation value. The level of priority outlined in the Operational Guidelines for the Implementation of the WHC to sites in Africa (along with the Pacific and Caribbean) to help close continental gaps (UNESCO 2019) could be applied for other representation measures, or to help make listing decisions between multiple sites in the same country or continent. The relatively low mnWHA inscription rate (~1.14 sites/year) suggests that representation gaps may take considerable time to close, but clear articulation of the representation components of interest and pathways to their achievement would signify progress for achieving global representation goals.

Closing some representation gaps to will require nominations and management to be reimagined. For example, marine wilderness inherently represents many of the criteria for natural outstanding universal value, making the WHC a promising mechanism for its conservation (Kormos et al. 2016; Allan et al. 2018). In fact, wilderness coverage could increase nearly eight-fold if just 15 of the tentative mnWHAs with the greatest marine wilderness area were inscribed. However, 90% of marine wilderness lies in offshore areas (Jones et al. 2018) and no mechanisms currently exist for inscribing WHAs beyond national jurisdictions (Abdulla et al. 2014). Recent WHC recommendations highlight the recognition of preserving such sites (UNESCO 2011), which is likely to become increasingly important as human pressures intensify (Eassom et al. 2016). Determining governance, monitoring and management responsibilities are only some of many challenges for offshore mnWHAs, yet partnering with existing organisations (e.g. International Seabed Authority, Food and Agriculture Organisation, International Maritime Organization) and global conservation agreements (e.g. Convention on Biological Diversity, UN Sustainable Development Goals) could help to define paths forward (Ardron et al. 2014; Laffoley & Freestone 2017).

The tentative list of mnWHAs provides several opportunities for increasing the uniqueness and representation of the existing World Heritage network. For example, to maximize representation of global marine ecoregions, just 10 tentative sites could add 15 ecoregions that are currently unrepresented in existing mnWHAs (Table 2b). Just six tentative sites could also collectively increase the representation of eight of the world’s rarest marine ecoregions (Table 2c). If prioritising greater representation of threatened species, the top ten tentative sites would result in an addition of 53 species (Table 2d). Importantly, the most complementary tentative sites to the existing mnWHA network change for each potential priority we explored. However, 15 sites perform well across multiple objectives and may represent efficient areas for reaching the objectives outlined here and the overarching goals of the WHC. Bonaire National Marine Park, for instance, contains seven unrepresented threatened species, a high threatened species density (>7 spp. km^−2^), one unrepresented ecoregion, and is experiencing the greatest human impacts—thus enhancing the representativeness across most metrics and, arguably, requiring the greatest additional protection to safeguard its natural assets. Importantly, these rankings assume all other site characteristics, including their outstanding universal value, to be equal. While this may not be the case, calculating these simple metrics can uncover potential trade-offs between objectives and provide a more holistic view of current and tentative WHAs.

Representation plays an important role in global biodiversity conservation (Margules & Pressey 2000) and international conservation agreements (e.g. Convention on Biological Diversity Aichi Target 11), and is often measured at the ecoregion scale (Butchart et al. 2015; Kuempel et al. 2016; Jantke et al. 2018). There is debate whether it is necessary for WHAs to achieve representation since outstanding universal value remains the primary standard for World Heritage listing (UNESCO 1995; IUCN 2004). While representation is generally discussed at the country or regional level and in relation to cultural sites (UNESCO 1994a), there is no clear definition of what level(s) of representation (e.g. country, ecoregion, species) are appropriate for natural areas or what constitutes achieving a representative WHA network. At the ecoregion scale, is not surprising that the current mnWHA network is not representative; this should not be expected given the relatively few existing mnWHA compared to 232 marine ecoregions. However, a baseline understanding of representation across levels is essential for future listing decisions to transparently and explicitly consider representation of the WHA network.

### Safeguarding natural assets and mitigating impacts

Roughly one out of every five existing mnWHAs is facing very high human impacts, with more than half of these mnWHAs experiencing relatively large increases in impacts (ΔCHI > 0.5). Close attention needs to be paid to these sites to ensure human impacts are urgently reduced and outstanding universal value is adequately maintained. Sites that experienced increasing impacts from 2008 to 2013 (77% of mnWHAs) should also be carefully evaluated so increasing trends can be neutralized or reversed. Over half of all tentative mnWHAs face high or very high impacts, but overall have experienced smaller increases in impact over time. On one hand, this could signify the urgent need to protect these sites from being lost; on the other, it could compromise the sites’ ability to meet the requirements of integrity and thus result in disqualification from listing. Currently, nominations and State of Conservation Reports largely rely on qualitative descriptions of impacts, and there appears to be no standardized reporting guidelines within the nomination scheme (UNESCO 2015). Quantitative evaluations are needed to assess the integrity of nominated sites and ensure effective management actions are implemented. These evaluations should be systematic to ensure that changes, threats, opportunities, and gaps within management can be compared and contrasted to provide a better synthesis of WHAs.

Our analysis of the relative contribution of land, marine and climate impacts can help guide decisions surrounding which management actions to undertake at a given site. For example, Rock Islands Southern Lagoon WHA in Palau had the highest overall impact in 2013. Like most sites, the majority of this impact is the result of climate change pressures. The global nature of climate change along with the WHC international platform and its financing mechanisms provide unique opportunities to help reduce and manage these impacts. The WHC clearly recognizes this need, having just completed an online consultation to assess the scale and risks of WHAs arising from climate change (UNESCO 2020), but rapid implementation is necessary. Some tentative sites with very high impacts are driven primarily by direct (ocean- and land-based) pressures, such as the Guérande salt marshes (73% of CHI), La Maddalena Archipelago and Bocche di Bonifacio Islands (64%), and Tayrona and Sierra Nevada de Santa Marta National Parks (61%), which can be locally managed. In Rock Islands Southern Lagoon, ocean-based impacts contribute more to CHI than land-based impacts, so focusing on fisheries and marine-based management efforts are critical. In areas where land-based pressures are more prominent (e.g. the Sundarbans, Guérande salt marshes, Kizilirmak Delta Wetland and Bird Sanctuary), efforts to reduce land-based pollution and habitat destruction could provide the most benefit. For example, stricter run-off laws were passed in Australia in efforts to keep the Great Barrier Reef WHA off of the endangered World Heritage list (McCutcheon 2019). Formal cost-benefit analyses and assessments of localized threats should be mandatory for management actions within these highly valuable areas.

Our study reveals significant gaps in the current mnWHA estate in terms of representation and management. However, there are several caveats to our analysis that should be considered when interpreting our results. First, the spatial boundaries for many tentative mnWHAs are largely unknown, which meant we had to base boundaries on existing protected areas or omit sites with too much ambiguity. While this is likely to over or underestimate representation and impact metrics across the mnWHA network, this is currently the best estimate available. Even using the best available data, we recognise there are major data deficiencies, specifically time lags and data gaps for important threatened species and human impacts. For example, the impact data did not include a measure of the threat from tourism, which may be the largest stressor facing many natural WHAs (Pedersen 2002; Hadwen et al. 2007), and several taxa were not included in metrics of species representation. Further, our analysis highlights broad patterns in impacts and representation to allow for global comparisons, but analyses at locally-relevant scales should be used for monitoring and reporting decisions within individual sites and countries.

## Conclusions

WHAs are the gold standard for protecting and managing the world’s most outstanding cultural and natural areas. The WHC has been successful in bringing international recognition and funding to these sites to help ensure their persistence. Many success stories exist, including the recovery of green turtle populations in Aldabra Atoll in the Seychelles and the eradication of invasive species from Macquarie Island in Australia (Casier & Douvere 2016). We applaud the success of these efforts, but call for clearly defined objectives (e.g., what is meant by representative, balanced and credible) and more rigorous, quantitative evaluations of current and tentative WHAs in meeting the goals outlined in the convention and subsequent 1994 Global Strategy. . Relatively simple metrics can be used to gain a more holistic understanding of the biodiversity benefits and potential impacts within WHAs, while more sophisticated analyses can help provide scientific basis for prioritizing the inscription of tentative sites and guiding cost-effective management actions. The global attention on WHAs and the myriad of supporting decision-making processes and infrastructure have the potential to produce rapid and lasting changes in how the world’s natural world heritage—and biodiversity in general— is monitored and conserved, while significantly contributing to an equitable, representative protected area network that transcends national boundaries.

## Supporting Information

**Text S1.** Data

**Text S2.** Analysis

**Table S1.** Existing Marine WHA summary (excel file)

**Table S2.** Tentative Marine WHA summary (excel file)

## Acknowledgements

We would like to thank Hugh Possingham for generously providing feedback on a previous version of the manuscript.

## Author contributions

Conceptualization, C.D.K., B.A.S., M.D.; Methodology, C.D.K., B.A.S., M.D.; Visualization, B.A.S.; Data Curation, B.A.S.; Writing – Original Draft, C.D.K., M.D.; Writing Review & Editing, C.D.K., B.A.S., M.D.

## Data Availability

https://github.com/BAlexSimmons/mnWHA_data

